# Harnessing microbes for crop production: TS201 enhances maize yield and reduces pest damage

**DOI:** 10.1101/2025.05.22.655610

**Authors:** Man P. Huynh, Khanh-Van Ho, Anne Z. Phillips, Amanda M Fox Ernwall, Dalton Ludwick, Deborah Finke, Zhentian Lei, Allison Jack, Bruce E. Hibbard, Natalie W. Breakfield

**Affiliations:** Division of Plant Science & Technology, University of Missouri, Columbia, Missouri, 65211, USA; Metabolomics Center, University of Missouri, Columbia, Missouri, 65211, USA; NewLeaf Symbiotics, St. Louis, Missouri, 63132, USA; Plant Genetics Research Unit, USDA-Agricultural Research Service, Columbia, Missouri, 65211, USA; Biological Control of Insects Research Laboratory, USDA-Agricultural Research Service, Columbia, Missouri, 65203, USA

## Abstract

Microbial technologies are increasingly adopted to improve sustainable agriculture amid escalating economic, regulatory, and ecological pressures, yet few, if any, are supported by mechanistic understanding that translates to real-world performance. Here, we report TS201, a U.S. EPA–registered bioinsecticide, composed of *Methylorubrum extorquens*, that enhances maize yield and resilience under pest pressure. Across seven years (2016–2022) of field trials at 22 U.S. locations, TS201 increased yield and reduced lodging. Large-scale evaluations at 81 sites in eight U.S. states (2023–2024) confirmed its agronomic benefit. Mechanistically, TS201 induced biosynthesis of methyl anthranilate, a volatile insect-repellent compound, and triggered pest avoidance of treated roots. These findings reveal a novel plant-microbe-insect interaction and establish a systems biology framework for harnessing microbial plant resilience to advance crop production.

## MAIN TEXT

Plant-associated microbiomes host diverse microbial taxa that often form symbiotic relationships with their host plants, shaping development and influencing resilience to environmental stressors (*1*). These interactions underpin a fast-growing class of agricultural biologicals enhancing crop productivity while reducing reliance on synthetic inputs (*2*). Despite their promise, most microbial products remain poorly characterized beyond proof-of-concept, and few, if any, are supported by mechanistic insights that translate to real-world agronomic outcomes.

This disconnection limits broader adoption to advance sustainable agriculture that feeds a growing population in an era of intensifying regulatory, ecological, and economic pressures (*3*).

Among beneficial microbes, pink-pigmented facultative methylotrophs (PPFMs) in the *Methylobacterium* and *Methylorubrum* genera are widespread in the phyllosphere and rhizosphere (*4, 5*). These single-carbon metabolizers influence crop physiology by modulating nutrient uptake, hormone balance, and immune signaling that can improve crop productivity (*6*). Here, we investigate a bacterial solution TS201, composed of *Methylorubrum extorquens* that enhances maize yield and mitigates pest damage evaluated under real-world field conditions.

### TS201: field discovery and performance

We conducted several field trials across the U.S. Midwest to evaluate various naturally occurring PPFMs as biological inoculants (biostimulants) to enhance maize performance. In one such trial under high pressure of corn rootworms (CRW, *Diabrotica* spp.), the most serious pests of maize (*7*), a subsequent wind event caused extensive lodging in control plots lacking PPFM treatment. Remarkably, plots treated with selective PPFM strains remained upright. Lodging is one of the most visible and economically consequential outcomes of CRW damage, as larval root feeding weakens plant stability and, under environmental stressors such as wind or rain, causes plants to lodge or topple, resulting in yield losses and harvest inefficiencies (*8, 9*). This observation prompted multi-year field trials to identify the most effective strain capable of boosting yield and mitigating CRW damage. Through this process, a PPFM species, *M. extorquens* strain NLS0042, designated TS201, was selected and further investigated and developed into a commercial product.

From 2016 to 2022, we performed extensive small-plot field trials at 22 locations across the U.S. Midwest, encompassing a range of low to high corn rootworm pressures, to evaluate the impact of TS201 on maize yield. All trial sites included locations where CRW pressure exceeded the economic injury threshold as indicated by nodal injury scores (NIS) of 0.25, a common measure of CRW damage (*10, 11*), in untreated control plots. Across these sites over seven years, TS201 significantly increased maize yield (220 kg/ha or 3.5 bu/A) and reduced CRW damage, as shown by decreased NIS values, indicating enhanced root protection under pest pressure (Table 1, table S1). Yield and pest suppression benefits were consistent across both conventional corn hybrids and those expressing insecticidal traits targeting CRW, including Bt toxins and lethal dsRNA constructs (table S2). TS201 also reduced lodging by 20% under moderate CRW pressure (1.2 NIS, from 59% to 39%) and by 36% under high pressure (2.7 NIS, from 65% to 29%). Based on its demonstrated biopesticidal efficacy, TS201 was registered in 2022 as a biological insecticide (TS201™, EPA Reg. No. 95699-2, Terrasym®, NewLeaf Symbiotics, St. Louis, MO) with the U.S. Environmental Protection Agency (EPA) under the Federal Insecticide, Fungicide, and Rodenticide Act (FIFRA) (*12*).

**Table 1.** TS201 improves maize yield and root protection in small-plot field trials under corn rootworm pressure. Field trials were conducted over seven growing seasons (2016-2022) at 22 locations across the U.S. Midwest. Data represents phenotypic responses in TS201 treatment relative to untreated control. Values summarize key agronomic and root protection traits observed in replicated small-plot trials. NIS: Nodal injury score.

| Metric |  | TS201 treatment<br>compared to control | P-value | Trialing scale |
| --- | --- | --- | --- | --- |
| Performance | Yield | +220 kg/ha | 0.0108 | 22 locations<br>over 7 years |
|  |  | 11,706 kg/ha (control) vs.<br>11,926 kg/ha (TS201) |  |  |
| Root damage | Nodal injury<br>score (NIS) | -0.11 NIS | 0.0007 | 22 locations<br>over 7 years |
|  |  | 1.34 (control) vs. 1.23<br>(TS201) |  |  |

Following U.S. EPA registration, we conducted large-scale on-farm evaluations of TS201 during the 2023 and 2024 growing seasons across 81 geographically and agronomically diverse environments in eight U.S. states (Fig. 2a). TS201 was applied in combination with Terrasym® 450, a PPFM biostimulant containing *Methylobacterium gregans* and evaluated under commercial-scale conditions using grower standard practices (GSP), which included a range of maize hybrids, management strategies, and pesticide applications as determined by each grower. Across all locations, the combined treatment with GSP resulted in a significant yield increase of +351 kg/ha (5.6 bu/A; *P* < 0.001) and achieved a 72% win rate over GSP alone (*P* < 0.001; Fig. 2b). Importantly, these trials were conducted regardless of actual CRW pressure, enabling assessment of agronomic performance and economic return under real-world agronomic conditions. These results demonstrate that TS201 consistently enhanced yield across diverse genotypes, soil microbiomes, and climatic conditions. The integration of TS201 into crop production systems complements sustainable integrated pest management (IPM) frameworks, which emphasize productivity, resilience, nutrient use efficiency, and reduced ecological footprint (*13*).

**Figure 1.**
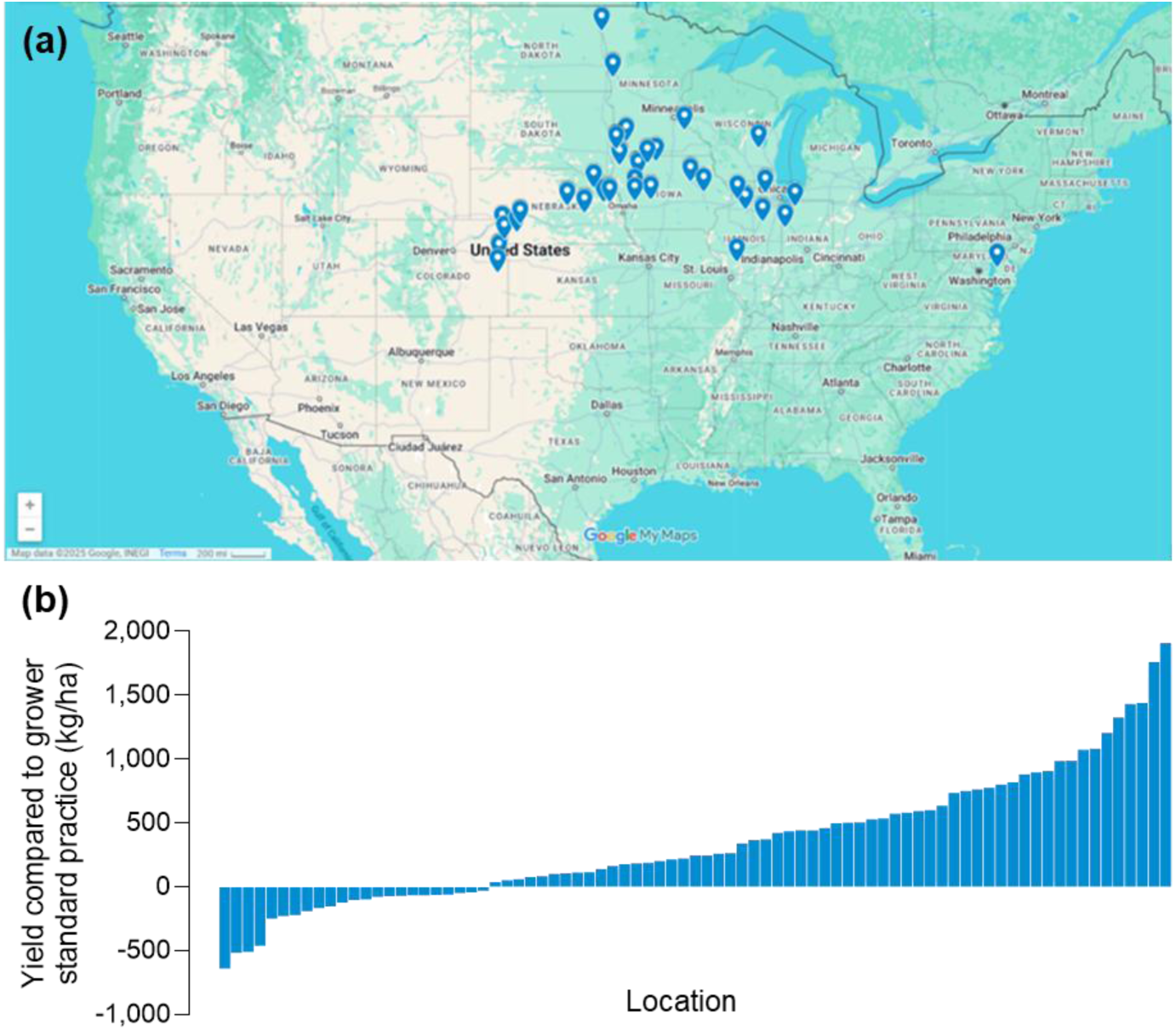
Large-scale on-farm evaluation of TS201 + Terrasym 450 in maize during 2023 and 2024 growing seasons across eight U.S. states. a) Geographic distribution of 81 commercial evaluation sites within eight U.S. states: Colorado, Iowa, Illinois, Indiana, Maryland, Minnesota, Nebraska, and Wisconsin. Trials were conducted regardless of corn rootworm pressure to assess real-world agronomic performance. Map background adapted from Google Maps (Map data ©2025 Google) with markers indicating county-level trial locations. b) Piano chart showing yield improvement of TS201 + Terrasym 450 + grower standard practice (GSP) as compared to GSP alone across large-scale on-farm trials (ranging from 4.05 to 40.5 ha per site, n=81).

**Figure 2.**
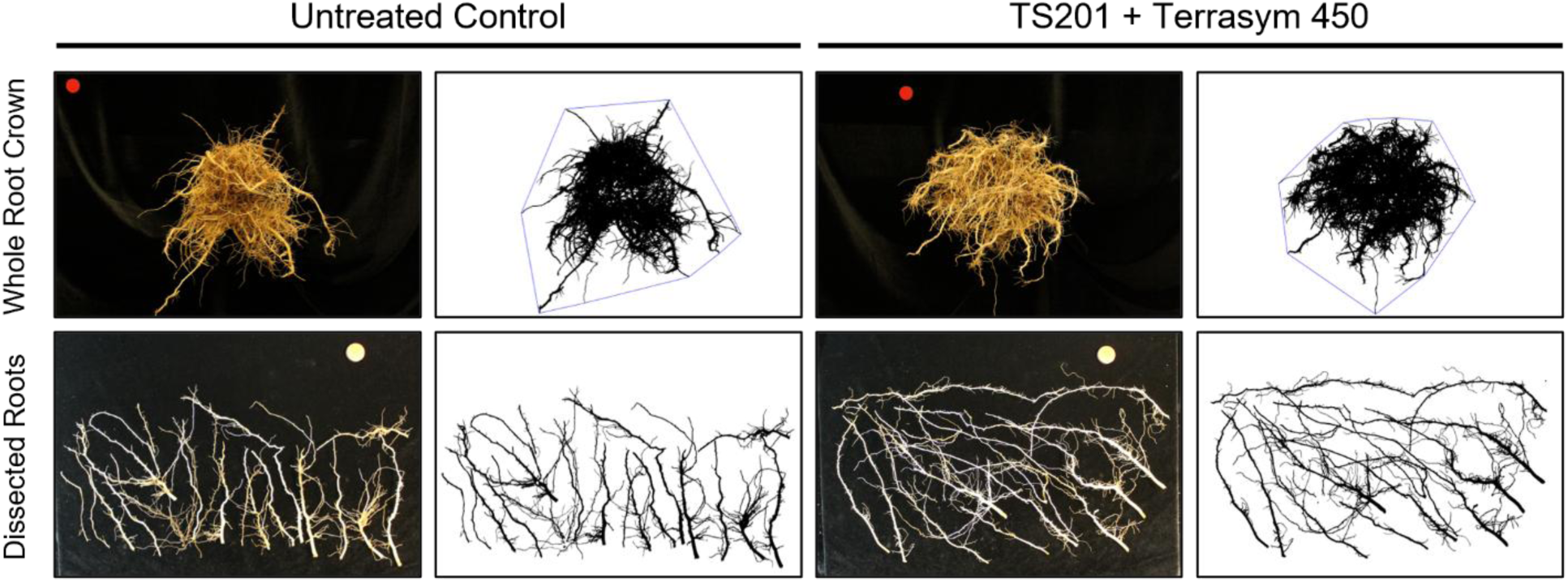
Enhanced maize root development observed under TS201 + Terrasym 450 treatment compared to untreated control. Root systems were collected from a small-plot field trial conducted in 2024 (10 plants per plot per replicate, four replicates per treatment). Following excavation and pressure washing, roots were imaged through a structured phenotyping workflow: intact root crowns imaged from below (top panel), then nodal segments were dissected and imaged (bottom panel). Images shown are representative of each treatment.

### Mechanisms of action

Prompted by prior observations of reduced CRW feeding damage and plant lodging, we conducted a small-plot field trial in 2024 to evaluate the effects of TS201 on maize root system architecture when applied in combination with Terrasym 450. The trial experienced in low but economically relevant CRW pressure, with a mean NIS of 0.26, slightly above the economic threshold of 0.25 (*10, 11*). Application of TS201 + Terrasym 450 resulted in consistent numerical increases in total root volume, surface area, cumulative root length, and root tip number relative to the untreated control, independent of visible feeding damage (Fig. 2, tables S3 & S4). The most pronounced and statistically significant enhancements were observed in nodal positions 1 through 3, where total root length increased by 25%, primarily driven by a 39% increase in fine root length (≤0.5 mm diameter) (table S4). These findings suggest that TS201 may provide early protection by preserving finer, actively growing roots that are preferentially targeted by CRW larvae (*7, 14*).

To investigate the molecular mechanisms by which TS201 influences maize physiology under CRW pressure, we performed RNA-Seq analysis on root tissues collected from a field trial in Illinois. Maize was planted and treated with TS201 in April, and root samples were harvested in mid-July, coinciding with root digs conducted to determine CRW feeding damage. RNA-Seq was conducted on TS201-treated root samples from plots that exhibited both reduced NIS and increased grain yield relative to untreated control plots (table S5). Differential gene expression analysis revealed five genes significantly upregulated in TS201-treated roots compared to controls (table S6), several of which are annotated as being involved in stress response and defense pathways based on the B73 maize reference genome. An additional eight genes were significantly downregulated (table S7). Notably, two of the upregulated genes were associated with the anthranilate biosynthetic pathway, prompting the hypothesis that TS201 treatment may induce the production of methyl anthranilate, a volatile compound previously documented to repel western corn rootworm (WCRW, *Diabrotica virgifera virgifera*) (*15*).

To determine whether TS201 elicits volatile defenses against herbivory, we performed two-choice olfactometer assays using WCRW (*16*), the most invasive and agriculturally significant pest within CRW species (*7, 17*). Although no behavioral differences were detected at 3 days post-treatment, by day 10, TS201-treated roots exhibited robust repellency to WCRW larvae compared to controls (Figs. 3a, c–f). This repellent effect was independent of prior root feeding and required a lag phase of more than three days post-application, suggesting that TS201 triggers volatile-mediated defense mechanisms through induced systemic resistance (ISR) (*18*) rather than damage-induced responses via systemic acquired resistance (SAR) (*19*). Notably, prior herbivory reduced larval avoidance (Figs. 3b, 3e), indicating that feeding may modulate volatile release or impose a fitness cost in maize. ISR, mediated by jasmonic acid and ethylene pathways, primes rapid, energy-efficient defense without constitutive gene activation (*20*). Emerging evidence also indicates that ISR orchestrates systemic metabolic reprogramming, hormone crosstalk, and oxidative signaling to enhance plant resilience (*21*). Although PPFMs are known to promote growth and pathogen resistance via ISR (*22–24*), their role in mitigating insect herbivory has remained largely unexplored. Our results demonstrate, for the first time, that a PPFM species can elicit a robust ISR response in maize that deters an herbivorous pest.

**Figure 3.**
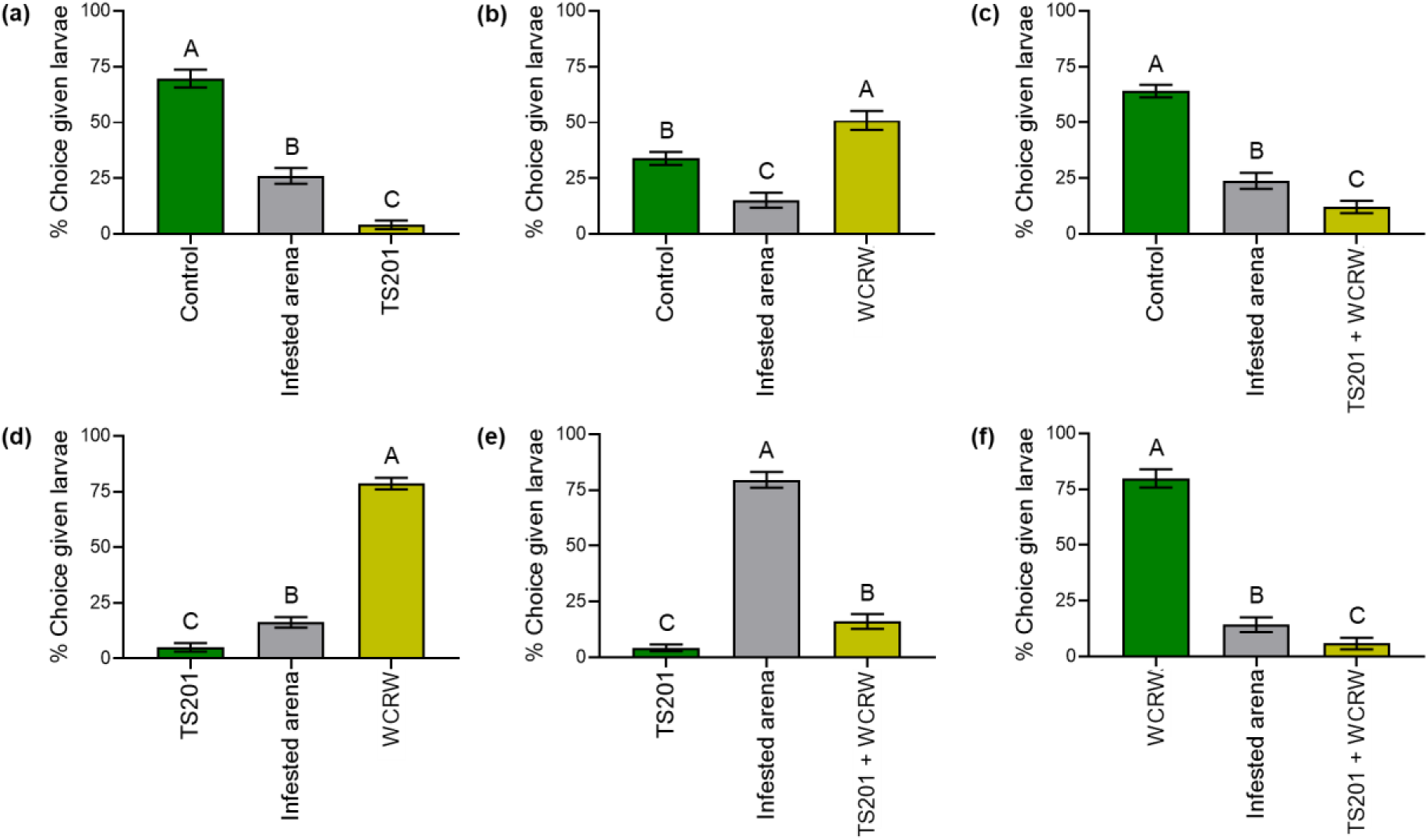
Percentage of larvae approaching experimental treatments in two-choice bioassays. Control: untreated seed without western corn rootworm (WCRW) infestation; TS201: TS201-treated seed without WCRW infestation; WCRW: untreated seed with WCRW infestation; TS201 + WCRW: TS201-treated seed with WCRW infestation. Bars with different letters are significantly different (*P* < 0.05). Mean ± SEM (n=12 independent choice assays).

To identify volatiles associated with WCRW repellency, we analyzed VOCs from TS201- treated and control plants using gas chromatography–quadrupole time-of-flight mass spectrometry (GC-QTOF-MS). Of the 18 VOCs detected (tables S8 & S9), methyl anthranilate was significantly induced by TS201 treatment (Fig. 4). Its relative abundance was markedly higher in TS201-treated plants, while the remaining 17 VOCs showed only minor or non-significant changes (Fig. 4). VOC levels were normalized to plant biomass, confirming that increased methyl anthranilate production was not due to plant size. Notably, this induction occurred independently of WCRW herbivory, with elevated emissions observed even in the absence of insect feeding. This emission pattern corresponded with the insect choice assays conducted on the same set of plants, where significantly fewer WCRW larvae were recovered from TS201-treated plants (fig. S1). Assays with purified methyl anthranilate further validated its strong, dose-dependent repellency against WCRW, with no significant repellency observed in untreated or solvent controls (fig. S2).

**Figure 4.**
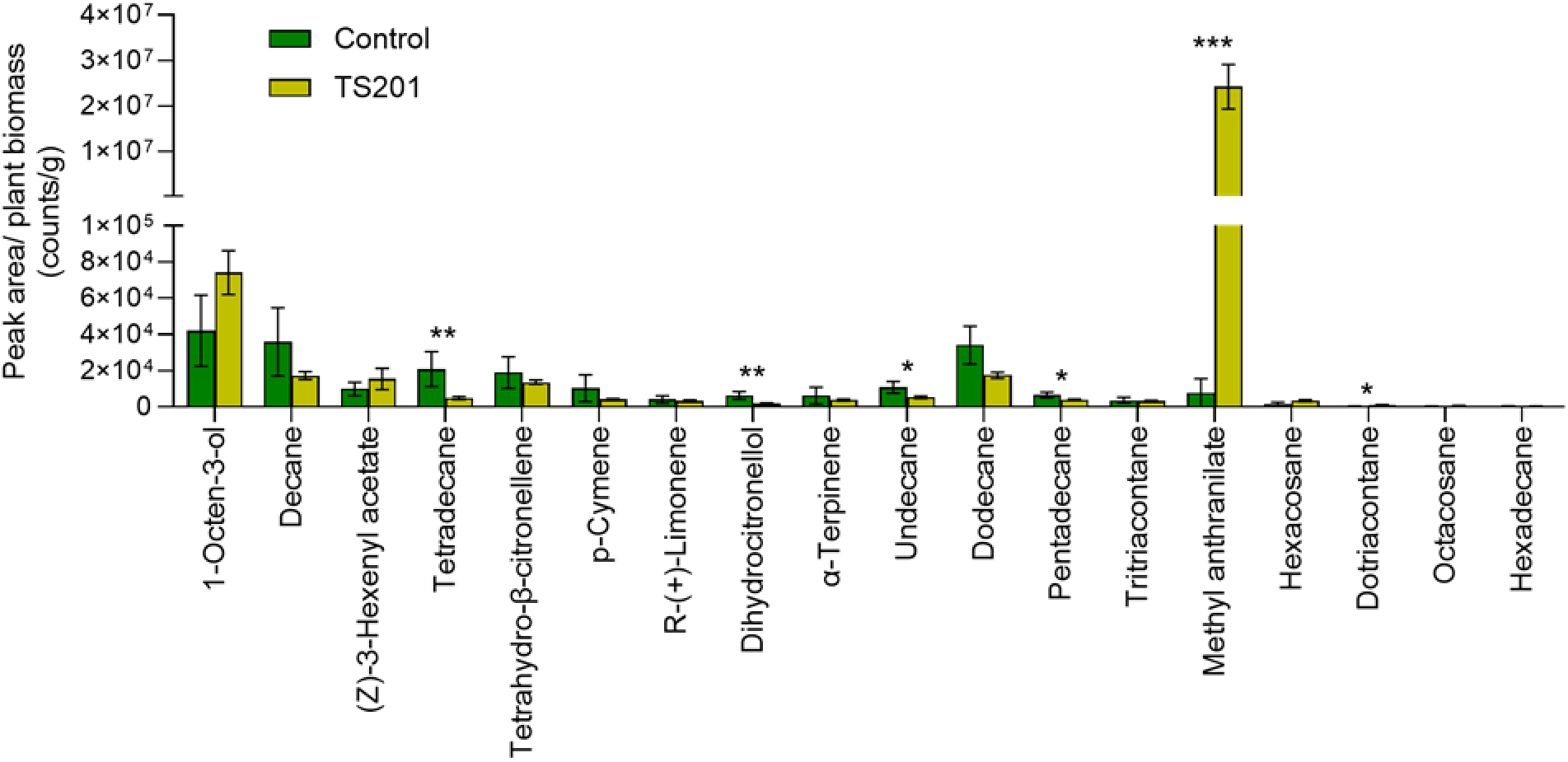
TS201 induces methyl anthranilate emission in maize. A list of all detected volatiles is available at Supplementary Table 7. Mean ± SEM (n ≥ 4). Untransformed data were presented, whereas analyses were performed with log10 transformed data. Signficant differences between the control and treated plants are indicated by *(*P* < 0.05), **(*P*<0.01), and ***(*P*<0.001).

Our results provide compelling evidence that TS201 activates the anthranilate biosynthetic pathway, leading to the production of methyl anthranilate, a volatile compound previously documented to repel WCRW (Fig. 5) (*15*). This mechanism represents a major advancement in IPM strategies for CRW, a pest complex that has evolved resistance to nearly all existing control strategies (*25–27*). Unlike conventional insecticides and Bt traits that impose direct lethal pressure and drive rapid resistance evolution, TS201 primes plant defenses through ISR, triggering endogenous pathways without direct toxicity, thereby minimizing selection pressure and potentially preserving non-target organisms (*28–30*). While biological control agents including *Heterorhabditis bacteriophora* and various biostimulants have shown variable efficacy and inconsistent field performance (*31–34*), TS201 consistently enhances maize protection across diverse agroecosystems, independent of pest pressure or environmental variability. These findings establish TS201 as a scalable, ecologically rational addition to sustainable CRW management, delivering strategic compatibility with current IPM frameworks and consistent efficacy across diverse geographic regions.

**Figure 5.**
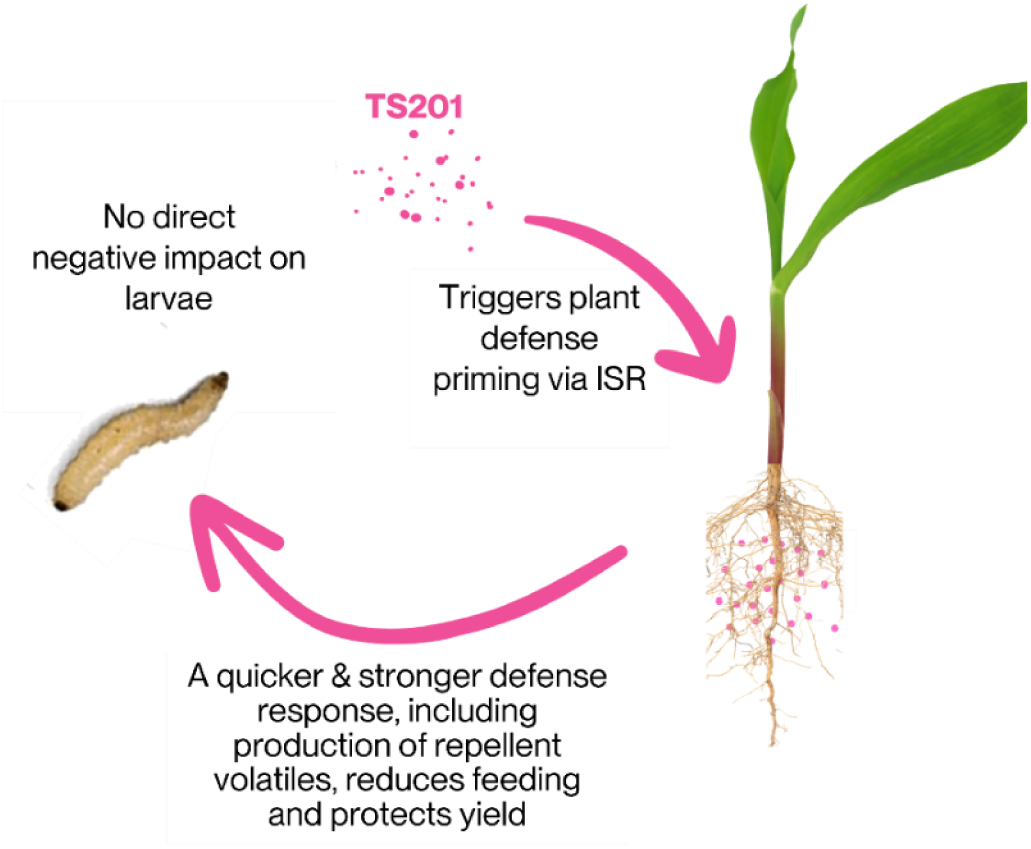
Schematic of TS201-induced enhancement of maize defense against western corn rootworm.

Plant-associated microbiomes represent a promising avenue for advancing climate-resilient, pest-tolerant cropping systems. Microbe-induced priming may concurrently enhance yield and strengthen plant defense under real-world agronomic conditions. Bridging field performance with mechanistic insight could accelerate the development of microbial solutions that reengineer plant resilience through ecologically informed partnerships for sustainable agriculture.

## ACKNOWLEDGMENTS

The authors thank the many NewLeaf employees both past and present who have worked on this project over the years including Vanessa Pugliessa for making the summary figure, Dayna Collett for running small plot trials 2018-2022, Roger Bowman for trial work in 2023-2024 and Ashley Haddox for supporting root phenotyping in 2024. We would also like to thank our trialing partners and contract researchers, especially Jay Pershing who consulted on this project for several years providing his extensive expertise in corn rootworm management. Funding for this project was provided by NewLeaf Symbiotics. This article reports the results of research only. Mention of trade names or commercial products is solely for the purpose of providing specific information and does not imply recommendation or endorsement by the USDA or the University of Missouri. USDA is an equal opportunity provider and employer.

## Author contributions

Conceptualization: M.P.H., B.E.H., and N.W.B.

Methodology: M.P.H., B.E.H., A.J., and N.W.B.

Investigation: M.P.H., K-V.H., A.Z.P., A.F.E., D.C.L., A.J., and N.W.B.

Visualization: M.P.H., K-V.H., A.Z.P., and N.W.B.

Funding acquisition: M.P.H., D.F., B.E.H., and N.W.B. Project administration: M.P.H., B.E.H., and N.W.B. Supervision: M.P.H., B.E.H., and N.W.B.

Writing-original draft: M.P.H. and N.W.B. Writing-review & editing: all authors.

## Data and materials availability

All raw transcriptomic sequencing data has been deposited in the Sequence Read Archive database on NCBI with BioSample Accessions SAMN48399254, SAMN48399255, SAMN48399256, SAMN48399257, SAMN48399258, SAMN48399259 under BioProject accession number PRJNA1260573. All other pertinent data are found in the figures and tables. Request for data and additional information should be submitted to the corresponding author.

## MATERIALS AND METHODS

### Biological materials

The EPA registered biopesticide product TS201, containing *Methylorubrum extorquens* strain NLS0042 (2.0%, 1 × 10^9^ CFU/g) as an active ingredient, was provided by NewLeaf Symbiotics (St. Louis, MO). This bacterial strain was originally sourced from a soybean plant in Saint Louis County, MO. Maize hybrids (P1197, Pioneer, Johnston, IA) were purchased from local seed suppliers unless otherwise indicated.

Western corn rootworm (WCRW, *Diabrotica virgifera virgifera*) neonates were obtained from a non-diapausing colony maintained at the Plant Genetics Research Unit-USDA-ARS in Columbia, MO (*35, 36*). The eggs were obtained in Petri dishes consisting of eggs and soil and were incubated at 25 ± 1 °C in darkness in an incubator (Percival, Perry, IA). Neonates that hatched within 24 h were used for insect assays.

### Field experiments

From 2016 to 2022, a series of 22 small-plot field trials were conducted across multiple locations in the U.S. Midwest with documented histories of elevated WCRW and northern corn rootworm (NCRW, *D. barberi*) damage, capturing variability in climate, soil type, and pest pressure. TS201 was evaluated as a standalone treatment to establish proof of concept under diverse field environments. During this pre-registration phase, trials were conducted under USDA-APHIS jurisdiction, with testing limited to a maximum of 4.05 ha (10 acres) per year in compliance with U.S. federal guidelines while U.S. Environmental Protection Agency (EPA) registration was in process.

The small-plot field experiments were designed as a randomized complete block design (RCBD) with four row plots 12 m (40 ft) in length and 4 to 6 replicates per location. TS201 was applied via multiple delivery methods to evaluate its efficacy and consistency under agronomically relevant deployment applications, including in furrow (309 g/ha, 1 × 10^9^ CFU/g wettable powder), liquid seed treatment (1 × 10^6^ CFU/seed application rate, 1 × 10^9^ CFU/g wettable powder) or planter box application (0.5 oz/ unit seed or 14 g/ 8 × 10^4^ maize seed, 1 × 10^9^ CFU/g powder with seed lubricity agent). In each trial, the cooperating contract researcher selected the maize hybrid and agronomic management practices with base insecticide treatments specifically excluded, and these plots served as the untreated control. TS201 application was the sole treatment variable distinguishing the treated plots from the untreated control. In a subset of these trials, the same maize hybrids were evaluated with and without belowground corn rootworm (CRW) protection traits such as Bt traits, with TS201 application was the only difference from the untreated control. Trials were conducted under continuous maize cultivation (corn-on-corn rotations) with select sites incorporating squash trap crops to facilitate consistent CRW feeding pressure. Root damage was evaluated in mid-July to early August, corresponding with the observed emergence of adult CRW beetles on maize plants, using the Iowa State University NIS system. To determine the efficacy of TS201 under CRW pressure, only field trials in which untreated control plots exhibited an NIS score greater than 0.25, a common economic damage threshold, were analyzed. Grain yield data were collected as bu/A and converted to kg/ha using the conversion of 1 bushel as 25.4 kg (*37*).

In 2020, an on-farm experiment was carried out to evaluate the impact of TS201 on maize lodging under CRW pressure. The trial included nine replicates in distinct locations in IA, IL, MN, and CO. At each site, 0.1 ha (¼-acre) strips were assigned to compare the untreated control with an in-furrow application of TS201 (309 g/ha, 1 x 10^9^ CFU/g). TS201 was applied to a corn hybrid without CRW protection traits. Roots were sampled along two defined transects (37 m) in each treatment, 20 roots total per treatment, pressure washed and rated using the NIS system. Out of nine locations, two in IA exhibited larval root injury exceeding the economic threshold (NIS > 0.25) and were included in lodging analysis. Based on NIS values, trial sites were categorized into low (0.25–1), moderate (1–2), or high (2–3) levels of CRW pressure levels. Lodging was quantified as the percentage of lodged plants within two, 37 m predefined transects for each treatment.

Following EPA registration, field evaluations of TS201 shifted from small-plot studies to large-scale, on-farm evaluations under commercial production conditions during the 2023 and 2024 growing seasons. The large-scale field experiment included 81 replicates, each representing a maize field at different locations within eight U.S. states (Fig. 2a). Across 81 trials (≥ 4.05 ha each), conducted in collaboration with contract researchers and commercial growers, TS201 was applied in combination with Terrasym 450 as a planter box treatment (0.5 oz per unit of seed of each PPFM isolate at 1 x 10^9^ CFU/g with seed lubricity agent) as an addition to the grower standard practice (GSP) and evaluated in comparison to GSP alone for CRW management. GSP reflected real-world production systems incorporating diverse IPM strategies, including combinations of belowground CRW protection traits (e.g., Bt, RNAi), in-furrow insecticides, crop rotation, and prior-season aerial applications targeting adult beetles. Trials were primarily conducted in corn- on-corn systems in regions with a reported history of CRW pressure. Experimental designs included full-field side-by-sides, split-planter setups, and replicated strips ranging from 4.05 ha to over 40.5 ha. Win rate was determined by dividing the number of trials with a positive yield over GSP by the total number of trials and converting to a percentage. Yield data were collected using precision monitors or weigh wagons and converted from bu/A to kg/ha as described above. Data were expressed as differences relative to the GSP control.

In 2024, building on prior observations of reduced CRW feeding damage and plant lodging, a small-plot field trial was conducted to evaluate the impact of TS201 on maize root system architecture when applied in combination with Terrasym 450. The trial was designed as a RCBD with four replicates. To determine baseline CRW pressure, roots from untreated control plots were sampled, pressure washed, and scored using the NIS system. Three weeks later, 10 root crowns per replicate were collected from both TS201 and Terrasym 450 treated and control plots to assess root architecture under CRW pressure. Roots were imaged using a structured phenotyping workflow: intact crowns imaged from below in the transverse plane and dissected nodal segments imaged per individual crown to enable accurate imaging of total root length, root size classes, and numbers of root tips. Root phenotyping images were analyzed using RhizoVision Explorer v2.0.3 (*38*). All images included a size marker for normalization and to convert pixel dimensions to metric units (mm). Dissected roots were grouped by node: 1-3, 4, 5, and 6. No node 7 roots were present in this study. Yield data were not collected, as the trial employed destructive sampling for detailed root phenotyping.

### Transcriptomic analysis

Root tissue samples from untreated and TS201-treated maize plants (one sample per plot; six replicates per treatment) were collected from different RCBD plots in a field trial. Total RNA was extracted using RNeasy Plant Mini Kits (74104, Qiagen, Germantown, MD), treated with DNase using the RNase-Free DNase Set (Catalog #15200, Qiagen), and assessed for RNA quality using an Agilent 2100 Bioanalyzer with RNA 6000 Nano Kits (5067-1511, Agilent Technologies, Santa Clara, CA).

Root samples from TS201-treated plots that exhibited both increased yield and reduced nodal injury scores (NIS) compared to control plots (three plots total) were used for gene expression profiling using QuantSeq FWD 3′ mRNA-Seq using 75bp reads with V2 Illumina chemistry (Lexogen, Vienna, Austria). QuantSeq is a tag-based sequencing method that captures and sequences the 3′ end of polyadenylated transcripts, allowing accurate gene expression quantification via read counting (*39*). The B73 v.4 maize reference genome was used for read alignment. Differential gene expression analysis was performed using the BlueBee Genomics platform which utilizes DESeq2; genes were considered differentially expressed if the log_2_ (fold change) was greater than 1.5 and the adjusted *P*-value was less than 0.1 (*40*). All raw sequencing data have been deposited in the NCBI Sequence Read Archive database with BioProject accession number PRJNA1260573.

### Insect choice assays

Untreated maize seeds were rinsed and incubated in distilled water overnight in a Percival incubator maintained at 25 ± 1°C to promote germination. After incubation, seeds were dried out and treated with TS201 at a concentration of 1 × 10⁶ CFU/seed, or with sterile distilled water (0.01 mL/seed, untreated seed). Subsequently, 3 kernels were planted per 50-mL conical tube and grown in the Percival incubator for 10 days. Plants were watered with 2 mL of distilled water every other day. The experiment included four treatment groups: untreated seed without WCRW infestation (control), serving as the control, TS201-treated seed without WCRW infestation, untreated seed with WCRW infestation, and TS201-treated seed with WCRW infestation. To allow for ISR priming to occur, seeds were allowed to germinate and grow for 7 days prior to insect infestation. Insect feeding was designed for 3-days to allow the potentiated ISR mediated-defense response to occur, if required. On day 7, 8 WCRW neonates (<24 h old) were transferred to plants assigned to infestation treatments. On day 10, plants were removed from the soil using a fine paintbrush and used for insect choice assays.

Insect choice assays were conducted using a modified olfactometer system adapted from Branson and Ortman (*41*). The olfactometer consisted of three 50 × 15 mm Petri dishes connected in a linear arrangement via 12 mm segments of Teflon tubing (8 mm I.D. × 10 mm O.D.). The bioassays were designed as a completely randomized experiment. Ten WCRW neonates were introduced into the central arena, while treatment and control plants were randomly assigned to the terminal arenas at either end of the olfactometer. The experiment was conducted in complete darkness in the Percival incubator, and larval positions were recorded after 30 minutes to assess behavioral responses. The experiment was run in three cohorts at different times, with each cohort consisting of four replicates, for a total of 12 replicates. Each replicate used unexposed larvae and freshly prepared plant seedlings.

Larval behavioral response was quantified by calculating the proportion of larvae that responded (PLR), using the formula: PLR = Nr / Nt, where *Nr* is the number of larvae found in each zone (treatment side, control side, or infested center) at the end of the assay (30 min), and *Nt* is the total number of larvae introduced.

### GC-MS analysis

Volatile organic compounds from control and TS201-treated maize plants were collected using a modified headspace sampling system based on the method described previously (*42*). Briefly, maize seeds were treated with TS201 or sterile water, and the treated seeds were planted without WCRW infestation in 50-mL conical tubes as previously described. At 10 days post-germination, seedlings were gently removed from the soil using a fine paintbrush. Maize seedlings were transferred into a 50-mL collection chamber (5 seedlings per chamber) fitted with Teflon tubing at both ends. One end of each chamber was connected to a flow meter (B07B6HQDDW, Wuhan, China), which was linked to a Whatman^®^ carbon/HEPA filter capsule (WHA67041500, Sigma-Aldrich, St. Louis, MO) and a 6-arm air delivery pump. The opposite end was connected to a Tenax^®^ TA thermal desorption tube (35/60 mesh, 0.25 in. O.D. × 3.5 in. length, Tenax^®^ TA, 29747-U, Sigma-Aldrich) for VOC trapping. Air was drawn through the chamber at a flow rate of 2 mL/min for 90 minutes to collect volatiles. VOCs from blank chambers containing sterile water and purified methyl anthranilate (≥98% purity, W268224, Sigma-Aldrich) were used as negative and positive controls, respectively. Adsorbed compounds were eluted from the Tenax tubes with 2 mL of methanol (Optima™ LC/MS Grade, A456, Fisher Chemical, Waltham, MA) for GC-MS analysis. The volatile assay was conducted as a completely randomized experiment with more than 4 replicates. Each replicate consisted of freshly prepared, unexposed maize seedlings randomly assigned to individual chambers within the headspace sampling system.

Headspace volatiles were analyzed using an Agilent 7890B gas chromatograph (GC, Agilent Technologies, Santa Clara, CA) coupled to an Agilent 7250 quadrupole time-of-flight mass spectrometer (QTOF-MS). Sample injection was performed using a Gerstel multipurpose sampler. The GC-MS operating conditions were as follows: helium was used as the carrier gas at a constant flow rate of 1 mL/min; injection volume was 1 µL with a split ratio of 5:1; and the inlet temperature was maintained at 280 °C. The oven temperature program began at 50 °C (held for 2 minutes), ramped at 6 °C/min to 215 °C, then increased at 5 °C/min to 315 °C, where it was held for 12 minutes before cooling back to 50 °C. Separation was performed using an Agilent DB-5MS capillary column (60 m × 250 µm i.d. × 0.25 µm film thickness). The MS transfer line was maintained at 250 °C, and electron ionization (EI) was conducted at 70 eV. Mass spectra were acquired in full-scan mode over a mass range of 40-650 m/z.

Data processing, including deconvolution, alignment, and compound identification, was performed using MS-DIAL software, with spectral matching against an in-house mass spectral library. The peak areas of identified compounds were integrated using Agilent MassHunter Qualitative Analysis 10.0 and normalized to plant biomass (mg) to determine their relative concentrations.

### Insect choice assays with methyl anthranilate

To determine the behavioral response of WCRW larvae to methyl anthranilate (MA), insect choice assays were conducted using the olfactometer system described above. In each assay, purified methyl anthranilate (≥98% purity; Sigma-Aldrich) was applied to one side of the olfactometer arena at concentrations of 0 (solvent control, methanol), 0.1, 1, or 10 µg with and without untreated maize seedlings, while the opposite side contained untreated maize seedlings. Ten neonate WCRW larvae (<24 h old) were introduced into the central arena of the olfactometer, and larval distribution was recorded after 30 minutes in complete darkness. Each treatment was replicated at least five times. The proportion of larvae responding was calculated as described above to determine the repellent effect of methyl anthranilate at each concentration.

### Statistical analysis

The VOC data were analyzed using Student’s unpaired *t*-test (PROC TTEST, SAS 9.4, SAS Institute, Cary, NC). Compounds were considered significantly induced when P < 0.05 and log_10_(fold change) > 1. Other data (i.e., small-plot field trial, root phenotyping, large-scale field trial, insect choice data) were analyzed using analysis of variance (ANOVA) followed by pairwise comparisons of least squares means (LSMEANs) or Fisher’s least significant difference (LSD) for multiple comparisons. Analyses were conducted using a generalized linear mixed model in PROC GLIMMIX (SAS 9.4) or JMP v18.1.1 (JMP Statistical Discovery LLC, Cary, NC), with treatment specified as a fixed effect and year, location, block, or replication as random effects. For win rate analysis, trial-level outcomes were treated as binary responses (win = 1, loss = 0) and modeled using a binomial distribution. Normality and homogeneity of variance were evaluated using univariate analysis in PROC UNIVARIATE (SAS 9.4). Datasets that did not meet these assumptions were transformed prior to analysis: percentage variables were arcsine square-root transformed, and numeric variables were square-root transformed, to meet the corresponding distributional requirements.

## SUPPLEMENTARY FIGURES

**Figure S1.**
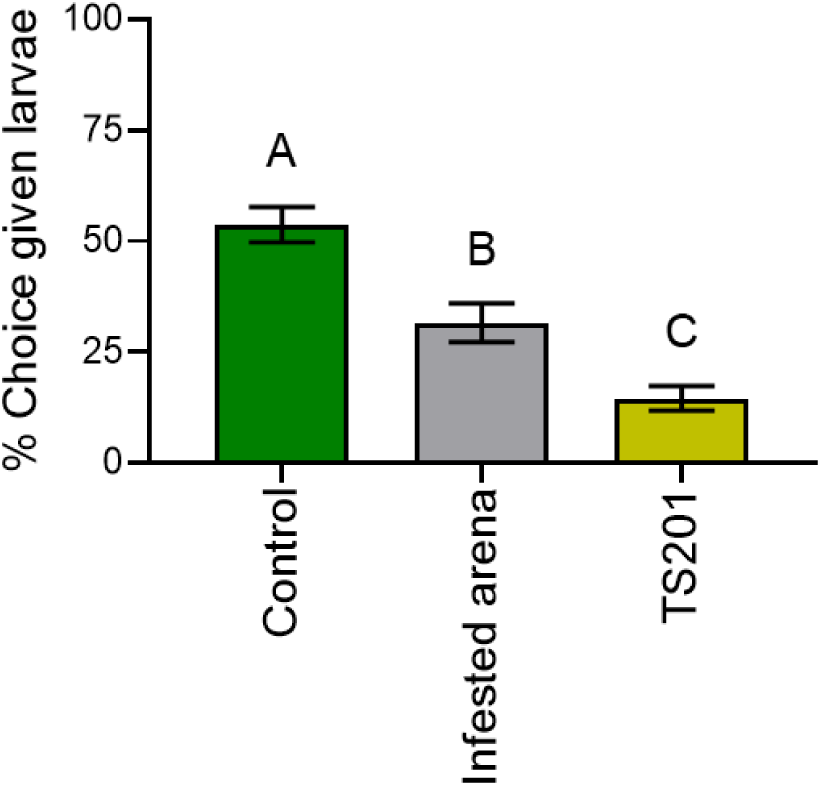
Percentage of larvae approaching experimental treatments in two-choice bioassays. Bars with different letters are significantly different (*P* < 0.05). Mean ± SEM (n=17 independent choice assays).

**Figure S2.**
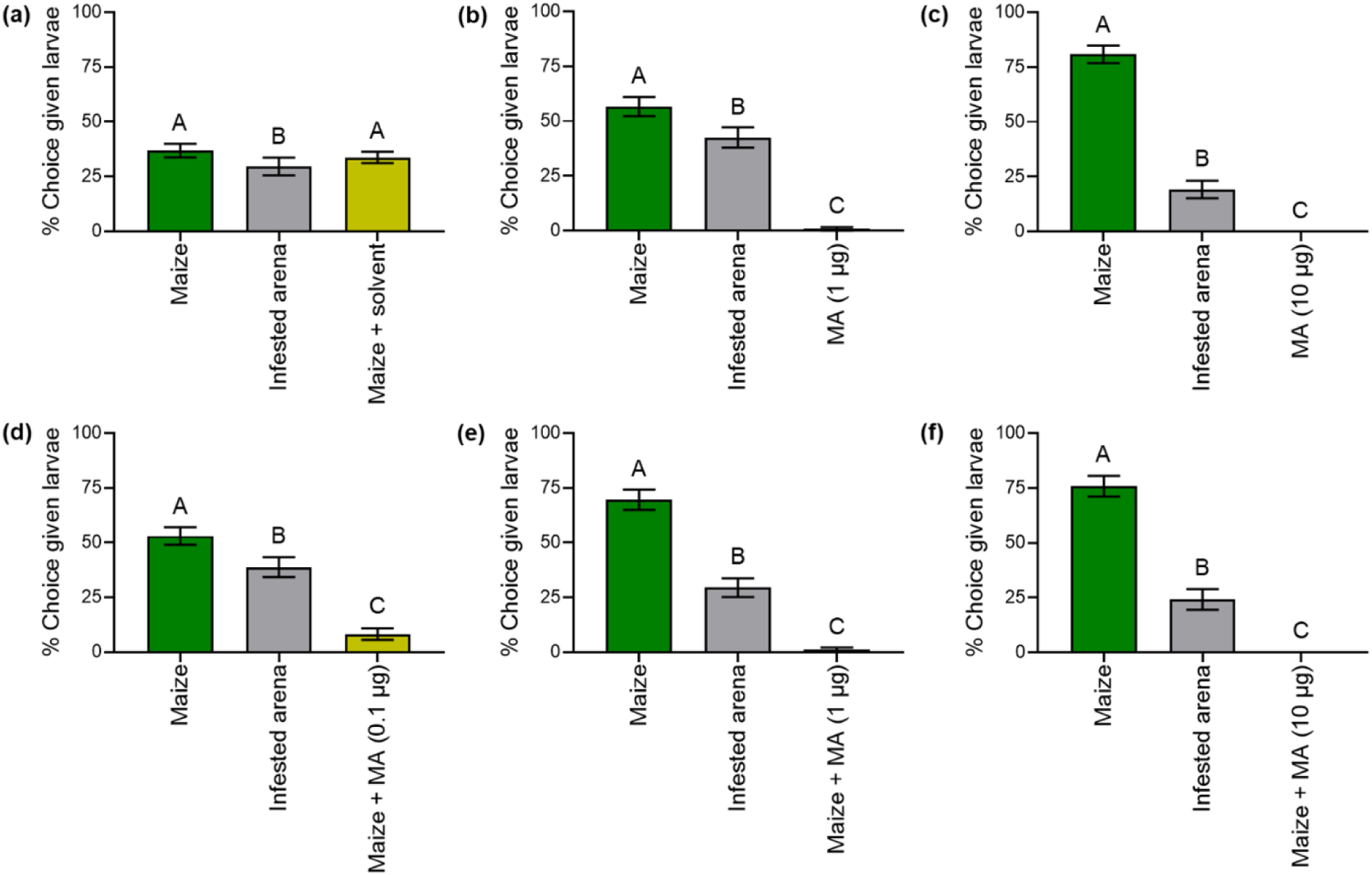
Percentage of larvae approaching experimental treatments in two-choice bioassays. Bars with different letters are significantly different (*P* < 0.05). Mean ± SEM (n=10 independent choice assays). MA: methyl anthranilate. Methyl anthranilate exhibited a strong, dose-dependent repellent effect against WCRW larvae, with few or no larvae found in its treatments. No significant difference was observed between untreated seeds and those with the solvent blank added.

## SUPPLEMENTARY TABLES

**Table S1.**
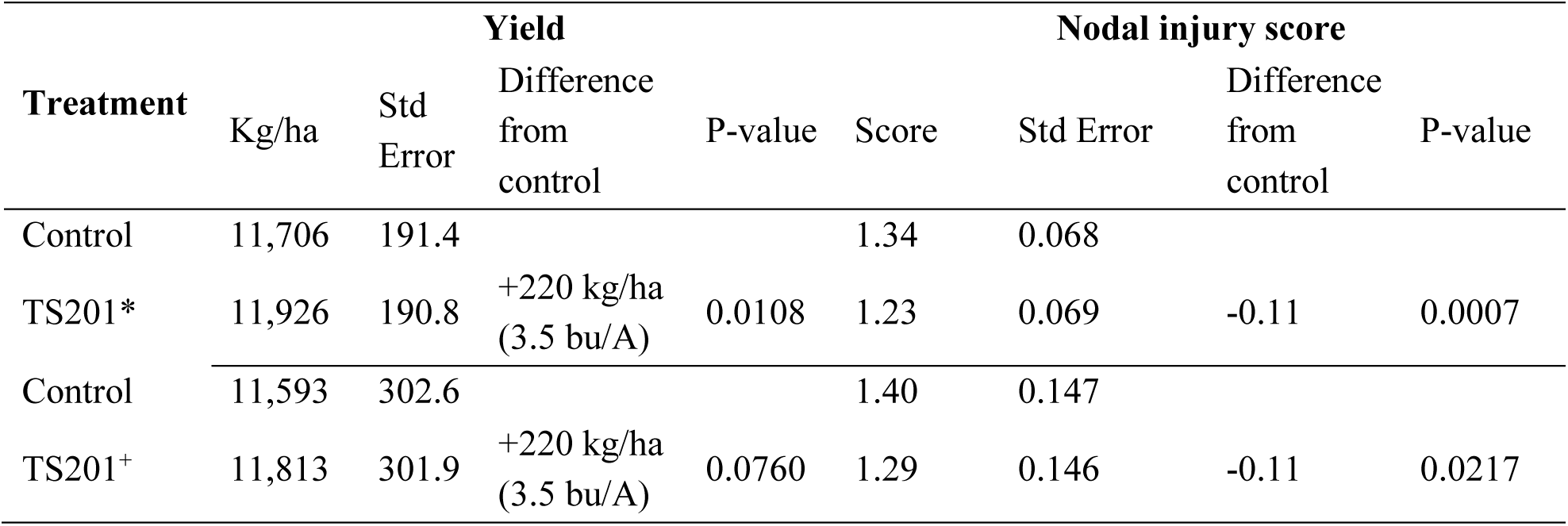
Effects of TS201 on agronomic performance and corn rootworm damage relative to untreated control (UTC). Field trials were conducted over seven growing seasons (2016-2022) at 22 locations across the U.S. Midwest. Data represent results from 22 small plot field trials. *Least square mean values when only TS201 and control were used in the linear mixed effects model; ^+^ Least square mean values when TS201, control, and all other treatments (i.e., other examined PPFMs, insecticides) in the trials were used in the linear mixed effects model.

**Table S2.**
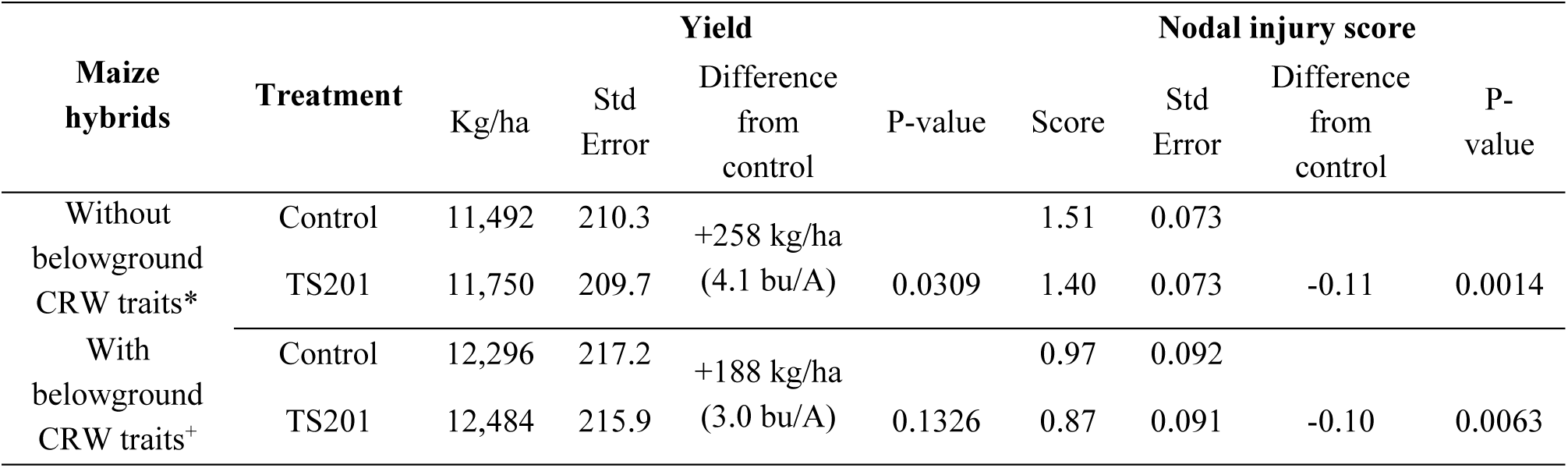
Effects of TS201 on agronomic performance and corn rootworm damage by corn rootworm (CRW) belowground traits relative to untreated control. *Data represent results from 22 small plot field trials in 2016-2022; ^+^Data represents results from 14 small plot trials over 2016- 2019 and 2021.

**Table S3.**
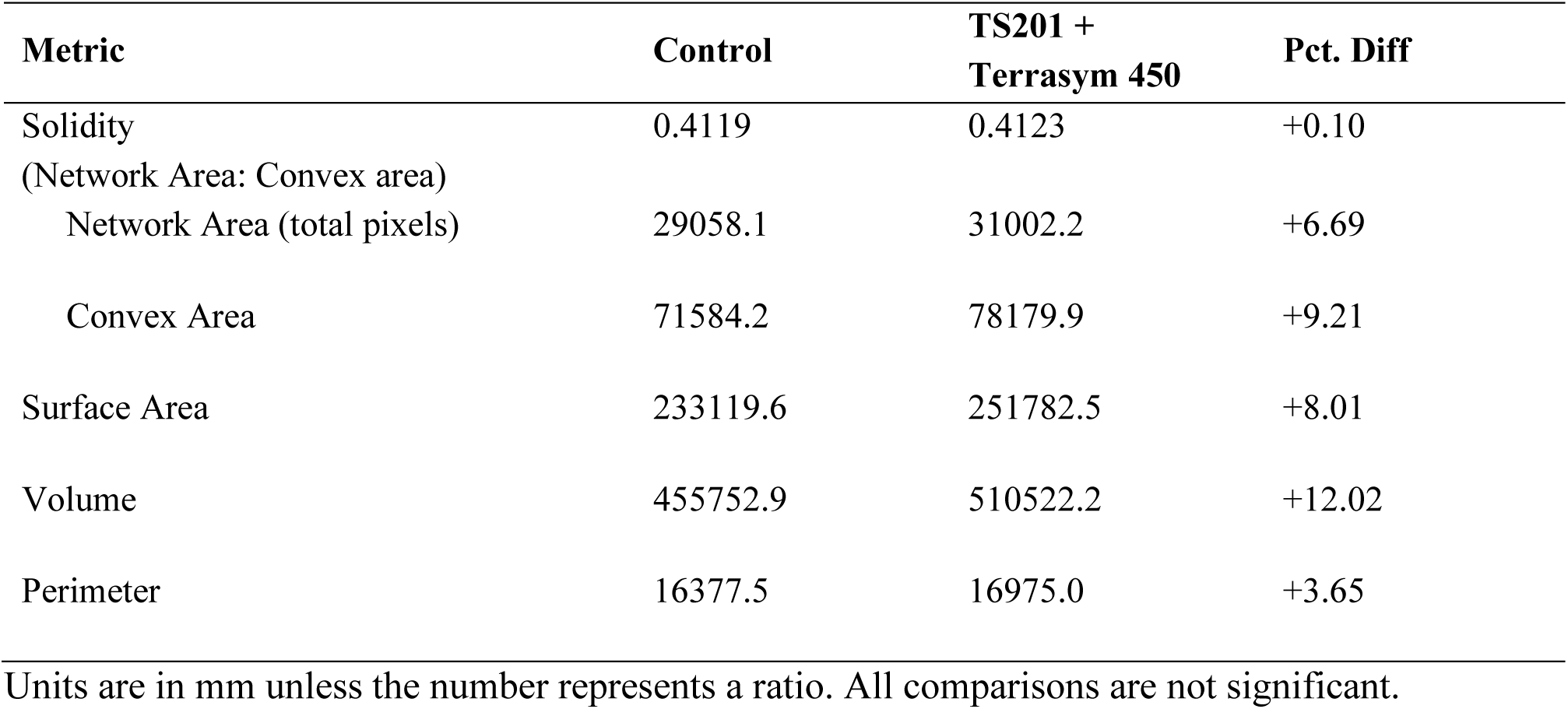
Full root crowns: Root phenotypes associated with the application of TS201 and Terrasym® 450 compared to an untreated control in a single location small plot trial with a baseline NIS of 0.26 in untreated control plots.

**Table S4.**
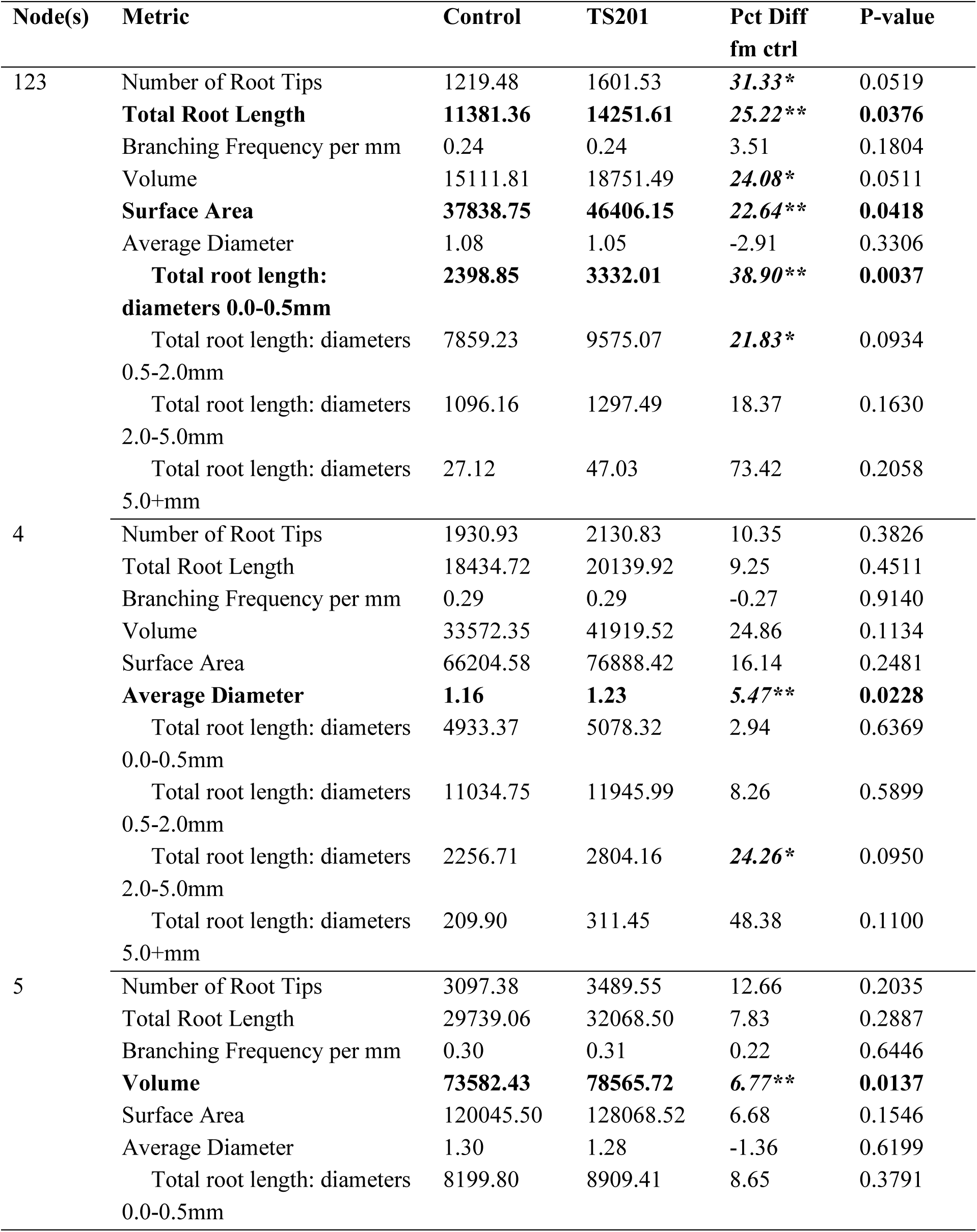
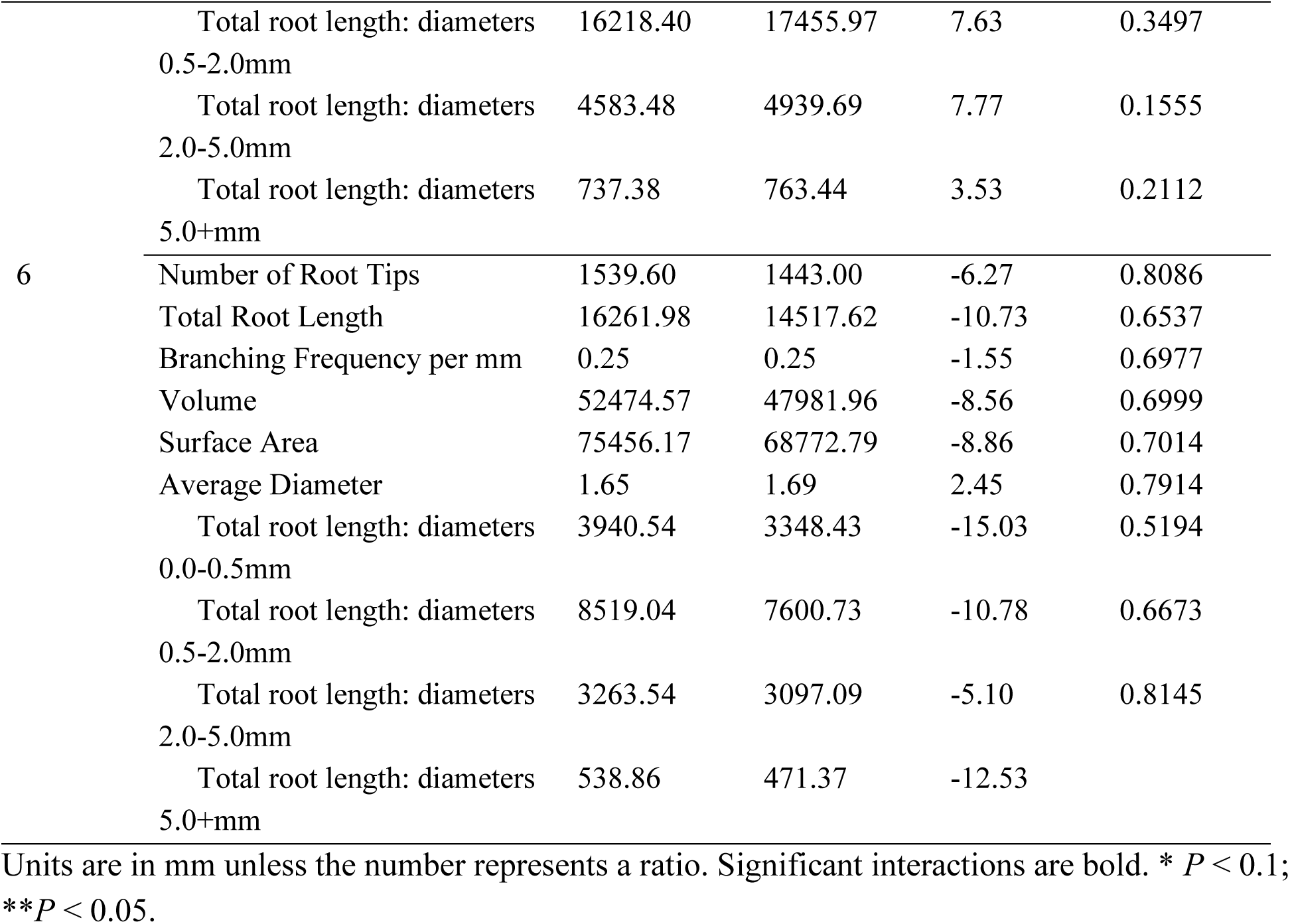
Dissected roots: Root phenotypes associated with the application of TS201 and Terrasym 450 compared to an untreated control in a single location small plot trial with a baseline NIS of 0.26 in the UTC plots.

**Table S5.** Field phenotypes of roots used in the RNAseq experiment.

| Treatment | Yield<br>(kg/ha) | Std<br>Error | Difference<br>from<br>control | P-value | Nodal<br>injury<br>score | Std<br>Error | Difference<br>from<br>control | P-value |
| --- | --- | --- | --- | --- | --- | --- | --- | --- |
| Control | 10,474 | 1479 |  |  | 0.35 | 0.08 |  |  |
| TS201 | 13,851 | 306 | +3,377 | 0.09 | 0.14 | 0.09 | -0.21 | 0.15 |

**Table S6.**
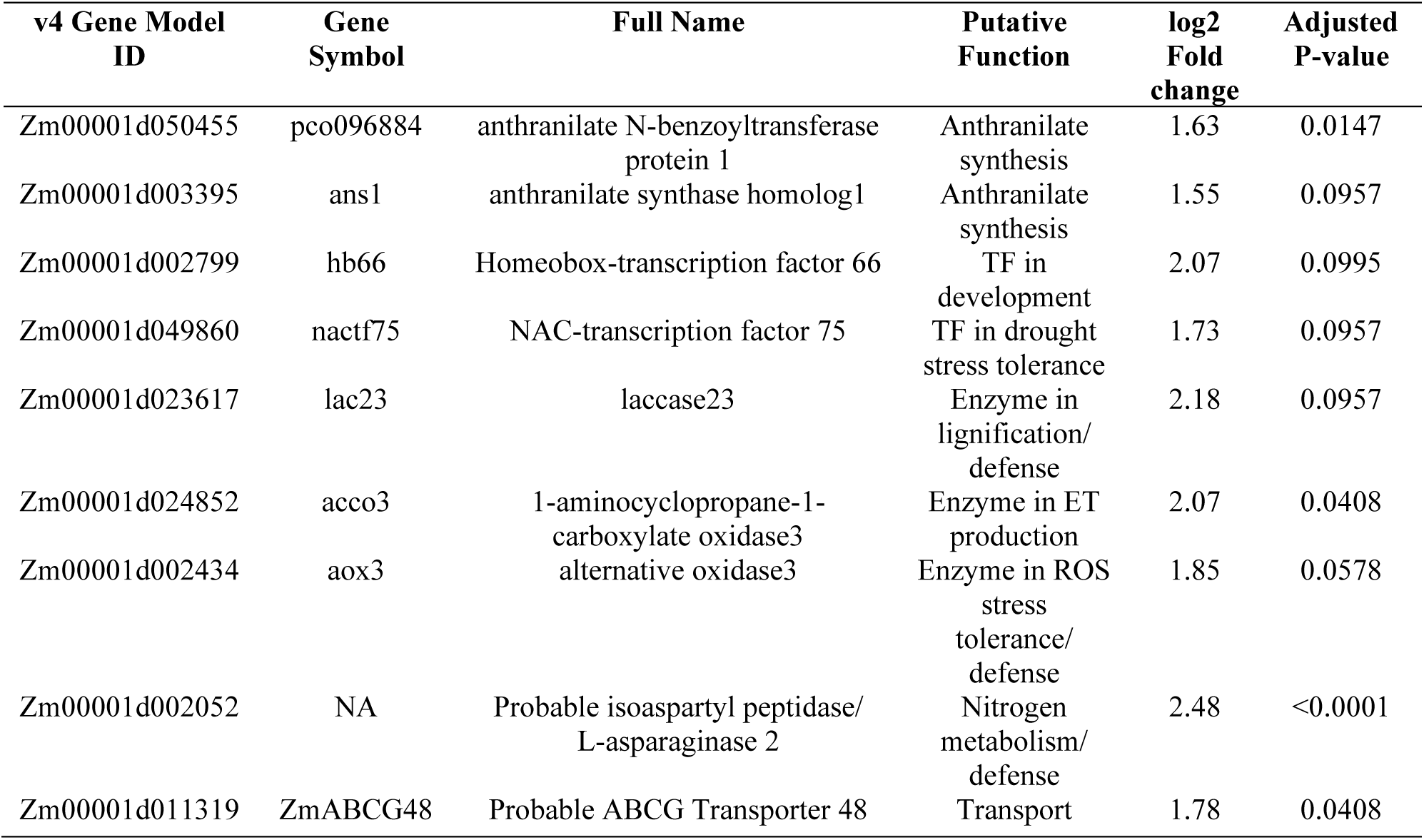
Up-regulated genes in TS201-treated maize roots relative to untreated control roots. TF= transcription factor, ROS=reactive oxygen species, ET=ethylene.

| v4 Gene Model<br>ID | Gene<br>Symbol | Full Name | Putative<br>Function | log2<br>Fold<br>change | Adjusted<br>P-value |
| --- | --- | --- | --- | --- | --- |
| Zm00001d050455 | pco096884 | anthranilate N-benzoyltransferase<br>protein 1 | Anthranilate<br>synthesis | 1.63 | 0.0147 |
| Zm00001d003395 | ans1 | anthranilate synthase homolog1 | Anthranilate<br>synthesis | 1.55 | 0.0957 |
| Zm00001d002799 | hb66 | Homeobox-transcription factor 66 | TF in<br>development | 2.07 | 0.0995 |
| Zm00001d049860 | nactf75 | NAC-transcription factor 75 | TF in drought<br>stress tolerance | 1.73 | 0.0957 |
| Zm00001d023617 | lac23 | laccase23 | Enzyme in<br>lignification/<br>defense | 2.18 | 0.0957 |
| Zm00001d024852 | acco3 | 1-aminocyclopropane-1-carboxylate oxidase3 | Enzyme in ET<br>production | 2.07 | 0.0408 |
| Zm00001d002434 | aox3 | alternative oxidase3 | Enzyme in ROS<br>stress<br>tolerance/<br>defense | 1.85 | 0.0578 |
| Zm00001d002052 | NA | Probable isoaspartyl peptidase/<br>L-asparaginase 2 | Nitrogen<br>metabolism/<br>defense | 2.48 | <0.0001 |
| Zm00001d011319 | ZmABCG48 | Probable ABCG Transporter 48 | Transport | 1.78 | 0.0408 |

**Table S7.** Down regulated differentially expressed genes in TS201-treated roots.

| v4 Gene Model ID | Gene<br>Symbol | Full Name | log2<br>Fold change | Adjusted<br>P-value |
| --- | --- | --- | --- | --- |
| Zm00001d015307 | NA | Uncharacterized protein | -2.23 | 0.006 |
| Zm00001d017454 | NA | NA | -2.08 | 0.058 |
| Zm00001d013259 | zmm15 | Zea mays MADS-box 15 | -2.07 | 0.058 |
| Zm00001d034045 | mads4 | MADS-transcription factor 4 | -2.04 | 0.048 |
| Zm00001d027763 | IDP1425 | plant-specific domain TIGR01615 family<br>protein | -2.03 | 0.007 |
| Zm00001d015092 | NA | NA | -1.89 | 0.096 |
| Zm00001d032457 | NA | NA | -1.86 | 0.041 |
| Zm00001d048137 | NA | NA | -1.86 | 0.015 |
| Zm00001d007045 | pco137705 | protein coding | -1.85 | 0.041 |
| Zm00001d029636 | cl945_1 | protein coding | -1.82 | 0.015 |
| Zm00001d035304 | limtf10 | LIM-transcription factor 10 | -1.75 | 0.006 |
| Zm00001d034621 | pco116267 | Protein trichome birefringence-like<br>32coding | -1.68 | 0.100 |
| Zm00001d043477 | cesa11 | cellulose synthase 11 | -1.65 | 0.096 |
| Zm00001d009506 | NA | NA | -1.58 | 0.100 |
| Zm00001d026287 | bnlg2190 | UDP-N-acetylglucosamine diphosphorylase,<br>bnlg2190 | -1.52 | 0.096 |
| Zm00001d043414 | NA | NA | -1.52 | 0.058 |
| Zm00001d000133 | NA | NA | -1.45 | 0.096 |
| Zm00001d019670 | mmp187 | kinesin-like protein KIN-4A | -1.44 | 0.096 |

**Table S8.** Volatile compounds annotated via GC-MS analysis.

| Compound | Formula | Retention<br>time | Quant Mass | log10<br>FoldChange | P-value |
| --- | --- | --- | --- | --- | --- |
| 1-Octen-3-ol | C<sub>8</sub>H<sub>16</sub>O | 9.415 | 57.03320782 | 1.13 | 0.3195 |
| Decane | C<sub>10</sub>H<sub>22</sub> | 9.885 | 57.06964660 | 0.96 | 0.4231 |
| (Z)-3-Hexenyl acetate | C<sub>8</sub>H<sub>14</sub>O<sub>2</sub> | 10.011 | 67.05367851 | 1.03 | 0.8210 |
| Tetradecane | C<sub>14</sub>H<sub>30</sub> | 10.191 | 57.06973394 | 0.88 | 0.0040 |
| Tetrahydro-β-citronellene | C<sub>10</sub>H<sub>20</sub> | 10.449 | 57.06957392 | 0.99 | 0.8237 |
| p-Cymene | C<sub>10</sub>H<sub>14</sub> | 10.719 | 119.0841019 | 0.98 | 0.8453 |
| R-(+)-Limonene | C<sub>10</sub>H<sub>16</sub> | 10.844 | 67.05382538 | 1.00 | 0.9776 |
| Dihydrocitronellol | C<sub>10</sub>H<sub>22</sub>O | 11.924 | 69.06966019 | 0.88 | 0.0018 |
| α-Terpinene | C<sub>10</sub>H<sub>16</sub> | 12.358 | 93.06890615 | 1.02 | 0.8063 |
| Undecane | C<sub>11</sub>H<sub>24</sub> | 12.552 | 57.06959788 | 0.93 | 0.0437 |
| Dodecane | C<sub>12</sub>H<sub>26</sub> | 15.37 | 57.06970382 | 0.95 | 0.0750 |
| Pentadecane | C<sub>15</sub>H<sub>32</sub> | 17.444 | 71.08510437 | 0.95 | 0.0493 |
| Tritriacontane | C<sub>33</sub>H<sub>68</sub> | 18.705 | 57.06973076 | 1.01 | 0.8138 |
| Methyl anthranilate | C<sub>8</sub>H<sub>9</sub>NO<sub>2</sub> | 19.502 | 92.02540970 | 2.93 | <0.0001 |
| Hexacosane | C<sub>26</sub>H<sub>54</sub> | 24.173 | 71.08522415 | 1.17 | 0.1235 |
| Dotriacontane | C<sub>32</sub>H<sub>66</sub> | 24.417 | 71.08522288 | 1.16 | 0.0125 |
| Octacosane | C<sub>28</sub>H<sub>58</sub> | 28.066 | 57.06988525 | 1.08 | 0.2463 |
| Hexedecane | C<sub>16</sub>H<sub>34</sub> | 29.027 | 71.08506775 | 0.97 | 0.5955 |

**Table S9.** Bioactivities of volatile compounds detected from maize.

| Compound | Pathway | Sub-pathway | Bioactivities | References |
| --- | --- | --- | --- | --- |
| 1-Octen-3-ol | Fatty acid metabolism | Oxylipin pathway | Induction of plant defense, insecticidal activity | (43); (44) |
| Decane | Fatty acid metabolism | Alkane biosynthesis | . | NA |
| (Z)-3-Hexenyl acetate | Fatty acid metabolism | Lipoxygenase pathway | Induction of plant defense, insect host and oviposition preference | (45); (46) |
| Tetradecane | Fatty acid metabolism | Alkane biosynthesis | Induction of plant defense, insect attractant | (47); (48) |
| Tetrahydro-β-citronellene | Terpenoid metabolism | Monoterpenoid biosynthesis | . | NA |
| p-Cymene | Terpenoid metabolism | Monoterpenoid biosynthesis | Insecticidal activity, insect repellent | (49); (50) |
| R-(+)-Limonene | Terpenoid metabolism | Monoterpenoid biosynthesis | Insect repellent | (51); (52) |
| Dihydrocitronellol | Monoterpene metabolism | Citronellol reduction | . | NA |
| α-Terpinene | Terpenoid metabolism | Monoterpenoid biosynthesis | Insecticidal activity | (53) |
| Undecane | Fatty acid metabolism | Alkane biosynthesis | Oviposition preference | (54) |
| Dodecane | Fatty acid metabolism | Alkane biosynthesis | Insect attractant, mating stimulant | (55); (56) |
| Pentadecane | Fatty acid metabolism | Alkane biosynthesis | Insect attractant | (57) |
| Tritriacontane | Fatty acid metabolism | Cuticular alkane biosynthesis | . | NA |
| Methyl anthranilate | Amino acid metabolism | Anthranilate biosynthesis (shikimate pathway) | Insect repellent | (58); (59) |
| Hexacosane | Fatty acid metabolism | Wax biosynthesis | . | NA |
| Dotriacontane | Fatty acid metabolism | Wax biosynthesis | . | NA |
| Octacosane | Fatty acid metabolism | Wax biosynthesis | . | NA |
| Hexedecane | Fatty acid metabolism | Alkane biosynthesis | . | NA |

